# Ensemble Processing and Synthetic Image Generation for Abnormally Shaped Nuclei Segmentation

**DOI:** 10.1101/2023.01.25.525536

**Authors:** Yue Han, Yang Lei, Viktor Shkolnikov, Daisy Xin, Alicia Auduong, Steven Barcelo, Jan Allebach, Edward J. Delp

**Affiliations:** Video and Image Processing Laboratory (VIPER), Purdue University; West Lafayette, Indiana, USA; HP Labs, HP Inc; Palo Alto, California, USA

**Keywords:** Nuclei Segmentation, Synthetic Data Augmentation, Ensemble, Machine Learning

## Abstract

Abnormalities in biological cell nuclei morphology are correlated with cell cycle stages, disease states, and various external stimuli. There have been many deep learning approaches that have described nuclei segmentation and analysis of nuclear morphology. One problem with many deep learning methods is acquiring large amounts of annotated nuclei data, which is generally expensive to obtain. In this paper, we propose a system to segment abnormally shaped nuclei with a limited amount of training data. We first generate specific shapes of synthetic nuclei groundtruth. We randomly sample these synthetic groundtruth images into training sets to train several Mask R-CNNs. We design an ensemble strategy to combine or fuse segmentation results from the Mask R-CNNs. We also design an oval nuclei removal by StarDist to reduce the false positives and improve the overall segmentation performance. Our experiments indicate that our method outperforms other methods in segmenting abnormally shaped nuclei.

## 1. INTRODUCTION

Quantitative analysis of nuclei morphology is important for the understanding of cell architecture. While most nuclei typically have an elliptical shape, deviations from this shape can arise in certain stages of the cell cycle, due to external stress, or in certain disease states. Some cell types also normally have non-elliptical nuclei (e.g. multi-lobed nuclei in neutrophils). Characterization of nuclei shapes therefore yields important information for many applications such as determining cell cycle stage, measuring cellular response to environmental stimuli, indicating genetic instability, and cancer diagnostics [1]. Traditional analysis of nuclei morphology requires manual assessment of a large number of microscopy images, which is laborious and time-consuming. Hence, image-based automated nuclei segmentation has been widely used to assist the researcher for the analysis of nuclei morphology.

Both traditional and deep learning image-based approaches have been used for automated nuclei segmentation. Traditional image segmentation methods such as Otsu [2] and Watershed [3] usually require manual parameter tuning and re-parameterization for new cell types and datasets to achieve an adequate performance [4]. Deep learning instance segmentation such as Mask R-CNN [5] and U-Net [6] provide very good results for nuclei segmentation after training on annotated nuclei groundtruth images [7]. To improve on these results, methods such as StarDist [8,9], Cellpose [10], DeepSynth [11], 3D Centroidnet [12], and NISNet3D [13] are specially designed for segmenting the nuclei in microscopy images. These methods have been developed for elliptical nuclei segmentation while approaches for segmenting non-elliptical nuclei are lacking.

Another challenge in nuclei segmentation is the difficulty in obtaining a large number of groundtruth images. One way to address this problem is by using data augmentation methods to create more training samples. Traditional data augmentation methods utilize linear and non-linear transformations including flipping, random cropping, color space transformations, and elastic deformations [14, 15]. However, these augmentation methods do not work well when the training data is limited and cannot solve the data imbalance problem within the dataset. With the development of Generative Adversarial Networks (GANs), generating synthetic images from GANs for data augmentation is widely used for various deep learning methods. SpCycleGAN was proposed to generate the synthetic microscopy images with the spatial constraint of the nuclei location [11, 12, 16].

Ensemble methods, where outputs of several segmentation methods are combined or fused, have proven to be useful in improving segmentation performance [17]. They are widely used in deep learning such as classification, regression, and segmentation [18]. The ensemble process can use different segmentation methods or the same segmentation method with different design parameters. In [19], different deep convolutional neural network architectures are used to improve the classification of cancer tumors. In [17], the results from multiple Mask R-CNNs are fused to segment the nuclei in different focal planes to reconstruct the 3D nuclei volume.

In this paper, we propose a system to obtain accurate segmentation masks with a specific type of abnormally shaped nuclei and a limited amount of groundtruth images. We first generate a specific abnormal shape of synthetic nuclei groundtruth images for training using SpCycleGAN. The shapes of synthetic nuclei will represent the special cases we encounter in our dataset. We train six different Mask R-CNNs based on sets of synthetic groundtruth images and then fuse the outputs using a type of non-maximum suppression with mask score averaging. We then use a pre-trained StarDist to reduce the false positives to obtain the segmentation. Our method can be used for many types of abnormally shaped nuclei by simply changing the shapes of the groundtruth mask generation. The proposed method outperforms several other methods for segmenting the abnormally shaped nuclei on our evaluation dataset.

## 2. PROPOSED METHOD

The block diagram of our proposed abnormally shaped nuclei segmentation system is shown in Figure 1. The system includes: (1) Synthetic groundtruth image generation for training, (2) 6 Mask R-CNNs with different architectures and training sets, (3) Ensemble fusion done by a modified version of Non-Maximum Suppression (NMS) with mask score averaging, (4) Pre-trained StarDist nuclei segmentation, and (5) Nuclei segmentation mask matching. The diagram shown on the left in Figure 1 is for the training parts of the system corresponding to the SpCycleGAN and the Mask R-CNNs. The inferencing is shown on the right in Figure 1. Note StarDist is a pre-trained system.

**Fig. 1:**
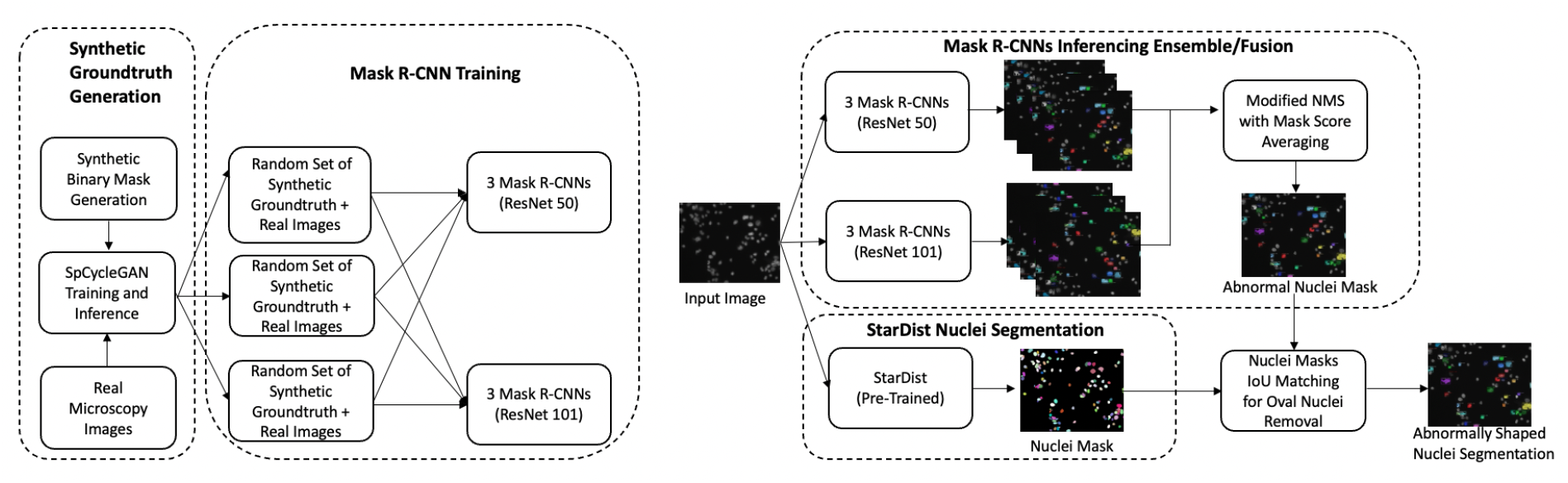
The block diagram of the proposed abnormally shaped nuclei segmentation system. The training system is on the left and the inferencing system is on the right.

Our goal is to locate a specific cell condition, which we are calling an unhealthy cell. Unhealthy cells are cells dying due to external stimuli during experiments. We desire to avoid fluorescent labeling of the cell membrane. Additionally, some cells are out of focus due to natural limitations in cell positioning in the imaging system, as well as limitations in the microscope, which makes cell boundary segmentation challenging. For these reasons, we choose to segment unhealthy cells’ nuclei for counting and localization of unhealthy cells. Figures 2a and 2b demonstrate two types of the unhealthy cell nuclei. Figure 2a is a nucleus which has irregular and small bumps on its surface (we shall call these type(i) nuclei). This specific shape is caused by the cells that are stressed and undergoing cell death, which will exhibit nuclei fragmentation and blebbing to look like small circular nuclei pieces budding off. Figure 2b is a nucleus that has an abnormal shape rather than oval (we call these type(ii) nuclei). Our goal is to segment only these abnormally shaped nuclei. Since type(i) nuclei are difficult to separate from typical oval nuclei and they occur infrequently, our system is designed to improve the performance of segmenting the type(i) nuclei.

**Fig. 2:**
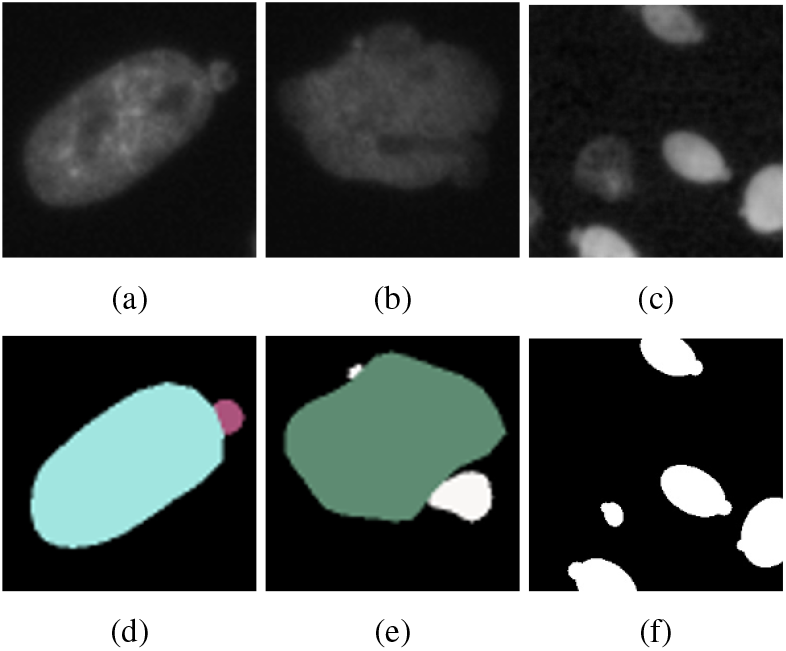
Unhealthy cell nuclei. The first row are images of nuclei and the second row are the segmentation masks. (a) and (b): type(i) and type(ii) nuclei respectively, (c): synthetic type(i) nuclei generated by SpCycleGAN. (d) and (e): StarDist segmentation masks of (a) and (b), respectively, (f): synthetic binary map from which (c) is generated.

### 2.1. Synthetic Groundtruth Generation

Because it is difficult to obtain ground truth images for this type of data, we use synthetically generated images. Synthetic groundtruth generation consists of synthetic binary map generation (segmentation groundtruth), SpCycleGAN training, and inferences (left side of Figure 1). To generate the synthetic binary maps of the type(i) nuclei, we first generate elliptical nuclei, then generate a ”circle bump” on the generated nucleus, where the center of the circle bump is located on the contour of the nuclei. The sizes and the numbers of nuclei and bumps are randomly chosen from ranges based on observing actual images. We add constraints to make sure the generated type(i) nuclei are not overlapping. We denote these synthetic binary maps as *I^ma^* and the original microscopy images as *I^ori^*

SpCycleGAN [16] is an extension of CycleGAN [20] with spatial constraints added to the loss function. CycleGAN combines 4 GANs, where *G* translates a binary segmentation mask to a microscopy image, *F* is for reverse translation of *G*, and two adversarial discriminators *D*_1_ and *D*_2_ are learned to make the two domain translations indistinguishable. SpCycleGAN adds one more network *H* to create a spatial constraint on the location of the nuclei and has the same architecture as *G*. The loss function of SpCycleGAN is shown in Equation 1.

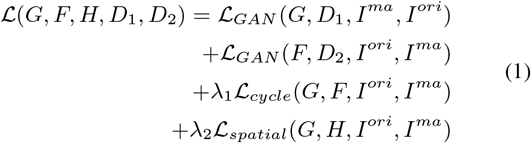

where 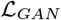 is the discriminators loss, *λ*_1_ and *λ*_2_ are the weight coefficients for the cycle consistency loss 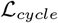 and the spatial loss *L_spatial_* proposed in SpCycleGAN. The spatial loss *L_spatial_* is defined in Equation 2.

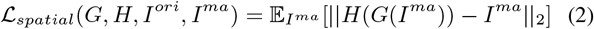

where ∥ · ∥_2_ is *L*_2_ is the norm. This minimizes the loss to ensure that the generated microscopy image has the nuclei in the correct position according to the binary segmentation map.

The SpCycleGAN is trained with unpaired real microscopy images *I^ori^* and synthetic binary segmentation maps *I^ma^*. It then generates synthetic microscopy images *I^syn^* corresponding to the *I^ma^*.

Figures 2c and 2f shows that SpCycleGAN can generate synthetic groundtruth images that well represent the shape and style of the type(i) nuclei.

### 2.2. Mask R-CNN Training

Mask R-CNN [5] is a two-stage instance segmentation network. The first stage of Mask R-CNN is to use a region proposal network (RPN) to generate bounding boxes indicating the potential objects, or we call them the regions of interest (ROIs). Then the second stage is used to classify and further refine the bounding box of the object, then provide the segmentation mask of the object inside the detected bounding box. The architecture of the Mask R-CNN is the feature pyramid network (FPN) [21]. FPN is designed to have lateral connections between each layer of the bottom-up and top-down convolutional layers to provide the predictions at multiple levels of the feature maps, thus maintaining strong semantically meaningful features at various resolution scales. We used the synthetic groundtruth type(i) nuclei images described above and a small number real ground truth images to train the 6 Mask R-CNNs.

### 2.3. Inference and Ensemble Processing

The generated synthetic abnormally shaped nuclei images do not perfectly represent the type of structures seen in real images. Since we only have a limited number of ground truth real images, training the Mask R-CNN with a large portion of synthetic images may cause the Mask R-CNN to learn the errors in the synthetic images. To reduce this, we randomly split the synthetic groundtruth images into different sets and combine them with the real images to create multiple training datasets for each of the 6 Mask R-CNNs. We then combine (or fuse) the outputs of the 6 Mask R-CNNs To fuse the output from each Mask R-CNN, we use a modified version of Non-Maximum Suppression (NMS) [22] with confidence score averaging to combine the segmentations from the individual Mask R-CNNs. We have *M* Mask R-CNNs, and the *i − th* Mask R-CNN’s output is denoted as 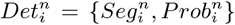, where 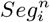 is a segmentation mask and 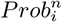 is its corresponding confidence score, *n* is the number of a detected nucleus and *i* ∈{1,…,*M*}. Our goal is to generate a refined final segmentation *Det^n^* based on 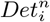. First we find the Intersection over Union (IoU) of the segmentation masks from each Mask R-CNN 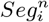. We then use Non-Maximum Suppression with a threshold of *τ* to construct the final segmentation mask *Seg^n^*. Unlike the original NMS which uses the highest confidence score to find the final confidence score of the mask, we use the average confidence scores to find the final confidence score *Prob^n^*. This ensures each Mask R-CNN’s output is contributing to the final segmentation. Equation 3 shows our modified NMS with mask score averaging.

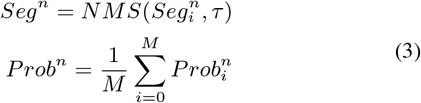

Where *NMS*(·, *τ*) is the Non-Maximum Suppression algorithm [22] with threshold IoU= *τ*.

### 2.4. Oval Nuclei Removal

Since the type(i) nuclei are very similar to typical elliptical nuclei if the bump is small, the existence of the false positive instances on the segmentation map still limits the performance. To resolve this issue without labeling more accurate data, we design an oval nuclei removal process using a pre-trained StarDist.

StarDist [9] is designed for segmenting nuclei, it uses a star-convex polygon to localize cell nuclei. During our experiments, we found that StarDist performs well for segmenting typical elliptical nuclei, but it tends to over-segment both type(i) and type(ii) nuclei into multiple instances, Figures 2d and 2e demonstrate this.

With StarDist’s characteristic of over-segmenting abnormally shaped nuclei, we use this characteristic as an indicator to remove typical oval nuclei from the results of the ensemble processing to reduce false positives. The block diagram of oval nuclei removal by StarDist is shown in Figure 3. We first count each closed contour from the StarDist segmentation as one mask then calculate the IoU between this mask and the results of the ensemble processing (Equation 4).

**Fig. 3:**
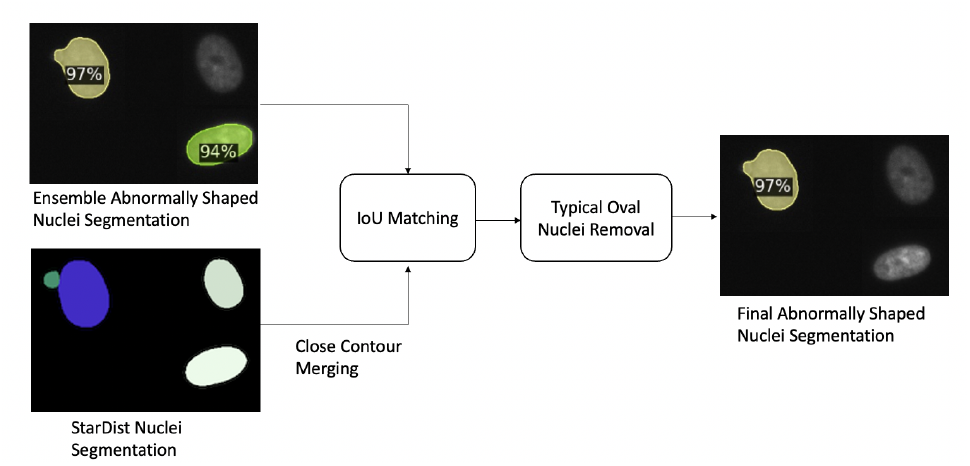
The block diagram of oval nuclei removal by StarDist.

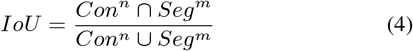

where *Con^n^* is the mask of *n −th* closed contour from StarDist, *Seg^m^* is the mask of *m −th* abnormally shaped nucleus from the ensemble processing.

If the IoU between *Con^n^* and *Seg^m^* exceeds the threshold *λ*, then these two masks will be considered as covering the same nucleus (IoU Matching). Once *Con^n^* contains only one segmented nucleus, the matched *Seg^m^* will be considered as an oval nucleus (false positive) and removed from the final segmentation.

## 3. EXPERIMENTS

We manually annotated type(i) and type(ii) nuclei from 50 fluorescence microscopy images (Figure 2). All the cells are CHO-K1 cells stained with ”Hoechst 33342 stain”. The images were captured at 200ms exposure with a DAPI filter cube on the microscope. The size of each image is 2758×2208 pixels. The dataset is divided into 30, 5, 15 images for training, validation, and testing. We manually selected 5 images with high nuclei density from the training dataset and randomly cropped 200 128×128 image patches for training the SpCycleGAN. We generated 200 128×128 synthetic groundtruth binary maps and along with 200 real image patches to form the unpaired training set to train a SpCycleGAN. The SpCycleGAN generated 600 synthetic type(i) nuclei groundtruth images based on 600 randomly generated binary masks. The images are randomly split into 3 sets evenly and combined with the same 30 real training images to form 3 training sets.

The 3 training sets are used for training 3 Mask R-CNNs with ResNet-50 and 3 Mask R-CNNs with RestNet-101 [23] architectures each. Each Mask R-CNN is trained on an Nvidia TITAN RTX GPU with a batch size of 16 and a base learning rate of 2.5*e^−^*^4^ for 25000 iterations. The IoU threshold for NMS for the ensemble processing is set to *τ* = 0.5. The pre-trained model for StarDist [9] is ”2D_ versatile_fluo” and the IoU matching threshold for oval nuclei removal is set to *λ* = 0.75.

### 3.1. Evaluation Metrics

We use the common metrics of precision, and recall to evaluate and compare the segmentation methods. A segmentation mask is considered as a true positive (TP) if the ground truth mask and it have an Intersection over Union (IoU) score greater than a given threshold. If a segmentation mask does not have a IoU score greater than a given threshold with any ground truth mask, it will be considered as a false positive (FP). A ground truth mask not being segmented will be considered as a false negative (FN). Precision and recall is defined as 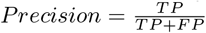 and 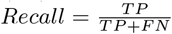.

### 3.2. Discussion

We compare our proposed method with one U-Net and one Mask R-CNN as shown in Table 1. Each closed contour on U-Net semantic segmentation will be considered as an instance mask of a nucleus. Both comparison methods are trained with the 30 real images and 600 synthetic type(i) nuclei images together. We also evaluate the ensemble segmentation without oval nuclei removal, denoted as Ensemble Only in Table 1. The confidence score threshold for the Mask R-CNN and U-Net is set to 0.8. The confidence score of the Ensemble Only is set to 0.6. The confidence score threshold for the proposed method is set to 0.4. The confidence score thresholds shown above are fine-tuned to produce the best result for each method on our validation dataset. Our proposed method achieves higher Precision and Recall than any other methods we compared to.

**Table 1:** Abnormally shaped nuclei segmentation performance

| Method | Precision | Recall |
| --- | --- | --- |
| Mask R-CNN [5] | 0.75 | 0.78 |
| U-Net [6] | 0.63 | 0.54 |
| Ensemble Only | 0.81 | 0.83 |
| Proposed Method | **0.84** | **0.88** |

Figure 4a shows the groundtruth of an abnormally shaped nuclei image. Figure 4b shows an example of U-Net segmentation, U-Net cannot differentiate the type(i) nuclei and typical elliptical nuclei well, resulting in several false positives and false negatives. Figure 4c shows the results for one Mask R-CNN and Figure 4d shows the results for Ensemble Only. The comparison between Figure 4c and Figure 4d demonstrates that the ensemble processing reduces the false negatives. The false negatives in Figure 4c are mostly type(i) nuclei, which confirms our assumption that the type(i) nuclei are the difficult case to be segmented due to their similar appearance to the typical elliptical nuclei. The ensemble processing does produce some false positives as shown in Figure 4d.

**Fig. 4:**
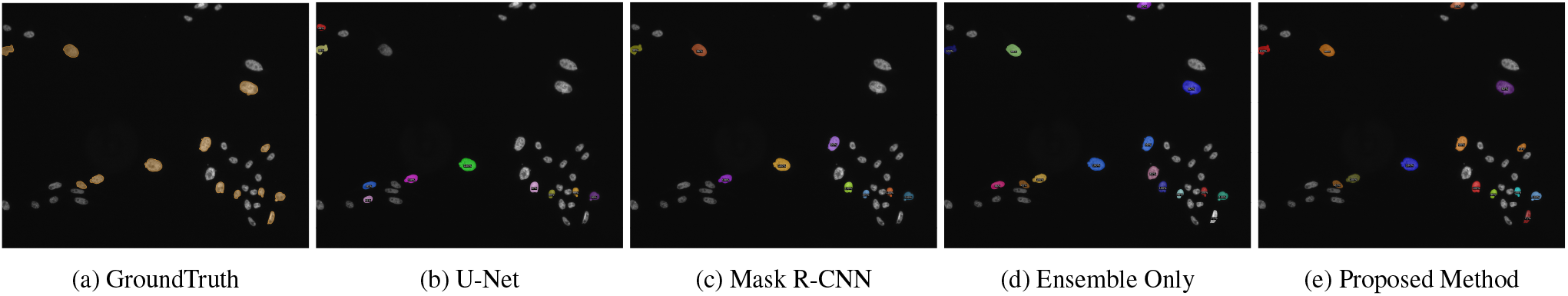
Segmentations for abnormally shaped unhealthy nuclei

By comparing Figure 4d and Figure 4e, we can see the oval nuclei removal done by StarDist removes most of false positives generated from the ensemble processing. Furthermore, with the help of oval nuclei removal, we are able to set a low confidence threshold and rely on it to reduce the false positives. With the low confidence threshold, the proposed method can successfully segment the difficult cases and reduce the false negative.

## 4. CONCLUSION AND FUTURE WORK

In this paper, we proposed a system to segment a specific type of abnormally shaped nuclei. We used a combination of synthetic groundtruth images and real ground truth images to train several Mask R-CNNs. During inferencing, the outputs of the R-CNNs were fused and compared with the outputs of a pre-trained StarDist to generate the final segmentation. This system performs well and can be generally used to segment various shapes of abnormal nuclei by simply changing the shape of the groundtruth generation mask. In the future, we want to investigate different loss functions which can better model the shape of abnormal nuclei and combined them into our segmentation method or synthetic nuclei generative model.

## 5. COMPLIANCE WITH ETHICAL STANDARDS

This is work is based on images obtained from commercially available cell line models for which no ethical approval was required.

## 6. ACKNOWLEDGEMENTS

This work was supported by HP Inc. Authors Yang Lei, Viktor Shkolnikov, Daisy Xin, Alicia Auduong, and Steven Barcelo are full-time employees of HP Inc. All other authors are with Purdue University and have no conflicts. Address all correspondence to Edward J. Delp at

